# Individualized network analysis reveals link between the gut microbiome, diet intervention and Gestational Diabetes Mellitus

**DOI:** 10.1101/2023.05.28.542631

**Authors:** Yimeng Liu, Guy Amit, Xiaolei Zhao, Na Wu, Daqing Li, Amir Bashan

## Abstract

Gestational Diabetes Mellitus (GDM), a serious complication during pregnancy which is defined by abnormal glucose regulation, is commonly treated by diabetic diet and lifestyle changes. While recent findings place the microbiome as a natural mediator between diet interventions and diverse disease states, its role in GDM is still unknown. Here, based on observation data from healthy pregnant control group and GDM patients, we developed a new network approach using patterns of co-abundance of microorganism to construct microbial networks that represent human-specific information about gut microbiota in different groups. By calculating network similarity in different groups, we analyze the gut microbiome from 27 GDM subjects collected before and after two weeks of diet therapy compared with 30 control subjects to identify the health condition of microbial community balance in GDM subjects. Although the microbial communities remain similar after the diet phase, we find that the structure of their inter-species co-abundance network is significantly altered, which is reflected in that the ecological balance of GDM patients was not "healthier" after the diet intervention. In addition, we devised a method for individualized network analysis of the microbiome, thereby a pattern is found that individuals with large deviations in microbial networks are usually accompanied by their abnormal glucose regulation. This approach may help the development of individualized diagnosis strategies and microbiome-based therapies in the future.

**Author Summary:** In this study, we aimed to investigate the role of the gut microbiome in gestational diabetes mellitus (GDM), a condition that affects pregnant women and is characterized by abnormal glucose regulation. Specifically, we asked whether and how the gut microbiome is affected by diabetic diet which is commonly used to treat GDM patients. We developed a new network approach to analyze patterns of co-abundance of microorganisms in the gut microbiota of GDM patients and healthy pregnant women. Our findings show that although the microbial communities remained similar after the diet phase, the structure of their inter-species co-abundance network was significantly altered, indicating that the ecological balance of GDM patients was not "healthier" after the diet intervention. Furthermore, we suggest that abnormal glucose regulation is associated with large network deviations, which could lead to the development of individualized microbiome-based therapies in the future. Our work highlights the importance of studying the microbiome from a network perspective to better understand the dynamic interactions among microorganisms in the community balance of the microbiome.

## Introduction

Gestational diabetes mellitus (GDM) refers to glucose intolerance that occurs or is first detected during pregnancy [1, 2]. GDM appears to be caused by the same physiological and genetic abnormalities as extra-gestational diabetes [3]. It is estimated to affect between 3-9% of pregnant women worldwide and is related with high rates of serious complications, for example shoulder dystocia and birth injuries, which includes bone fractures and nerve palsies[3, 4]. Babies born to mothers with GDM may have issues with persistent impaired glucose tolerance [5], subsequent obesity [6], and impaired intellectual achievements [7]. Furthermore, even after pregnancy, people with GDM may still have diabetes, which poses a high risk for people with GDM [8, 9]. Common treatments of GDM that aim to reverse hyperglycemia include lifestyle changes and insulin therapy [10]. Usually, lifestyle changes consist of diet intervention, exercise therapy and blood glucose self-monitoring. Insulin therapy is often used when lifestyle changes fail to control blood glucose levels or when complications arise with the fetus.

GDM is a typical metabolic disease that occurs during pregnancy, which may suffer from gut microbiome disorders. In fact, in recent years, many diseases, not only metabolic diseases, have been found to be closely related to flora disorders, including *Clostridium Difficile* infection (CDI) [11], colorectal cancer [12], dietary choline-induced atherosclerotic heart disease [13] and chronic diseases such as obesity [14]. In some cases, the change in the microbiome during a disease appears as an abnormal abundance of specific taxa [15]. However, a disease state can also be associated with a community-wide shift of the microbiome state (commonly evaluated in terms of PCA or *α*- and *β*-diversity measures). Such cases represent more general abnormalities that are linked to the interactions between the species and their ecological balance [16]. One of the most important factors that can influence microbiome composition is diet [17–19]. For example, the Mediterranean diet may benefit those with underlying conditions, such as obesity, blood lipids and inflammation [20]. Diet interventions may cause community-wide alterations of the microbiome, by affecting the ecological interactions via promoting or inhibiting microbial growth [21].

Thus it can be seen that understanding the underlying role of the microbiome of GDM patients is essential in two ways: On the one hand, an altered microbiome can affect their general health state. On the other hand, the individual composition of the microbiome may be related to the success of dietary interventions. Current studies of the microbiome in GDM patients have mainly focused on the abundance of specific microbial taxa [22, 23], but not on the global interaction structure of different taxa. These studies revealed that the proportions of certain microbes in GDM patients differ from those in healthy subjects [24, 25]. However, at the microbiome community level, there was no clearly difference in composition and structure of intestinal microbiome communities between GDM patients and healthy pregnant women at three different stages of pregnancy through PCoA and α-diversity analysis [1, 26]. Thus, it is still unclear whether alterations of the community structure of the microbiome play any part in the condition of GDM.

Considering traditional biological community methods were unsuccessful in fully revealing the changes of the microbiome community of GDM patients, here we wish to instead analyze the microbiome community from the network perspective of co-abundance. To unveiling the complex web of interactions in microbial communities, dynamic ecology and evolutionary processes which drives them are required to be understood [25]. Microbial community structure and their functions are complex because of their dynamic nature, variability in composition, their self-reproduce ability and self-organize ability. Therefore, this complexity can be well represented and modeled as a network[27, 28]. The interactions within microbial communities can be well analyzed through the network method. In addition, the network approach can also be used to analyze the role of microbial communities between disease and health [29], and thus can detect changes in the appropriate ecological balance, helping to identify the health of microbial community balance, and providing additional information about the underlying dynamics. In contrast to traditional microbiome analysis, which focuses on whether the abundance of individual species is within the normal range and detects abnormal abundances, the network analysis method focuses on the balance between species and detects abnormal ecological interactions.

Combined with the existing diet intervention strategies, in order to better understand the changes of microbiome in the course of GDM health evaluation and diet intervention, in this study, we propose a new approach which is based on individualized - and group-networks to analyze the changes in microbial communities between subjects with GDM and subjects without GDM (healthy), with and without diet intervention. The community structure and community balance are analyzed and compared with and without diet intervention by network similarity calculation. We reveal the effect of diet intervention on the microbiome of GDM patients and demonstrate the relationship between the network structure of the microbiome and the diagnosis of individual blood glucose. These findings provide support for evaluating the recovery of GDM patients, and contribute to the future personalized microbial-based medicine.

## Results

### Data collection and microbiome network analysis

The experimental design of this study consists of observing and collecting data from healthy pregnant women and GDM patients who received diet intervention before and after two weeks, that was not especially designed for this study. The oral glucose tolerance test (OGTT) is performed to all pregnant subjects at the first instance of stool collection. Women with abnormal blood glucose levels during the test were diagnosed with GDM and receive traditional diet intervention. Patients with GDM had their daily calorie intake tailored to their weight by a nutritionist (see Methods). Women with normal blood sugar levels were designated to the control, healthy group. In this observational study, diet intervention was applied by the hospital as part of the routine treatment for GDM patients, while, to mitigate ethical concerns, healthy pregnant women were not recommended any special diet intervention. Dietary intervention for patients with GDM is the suggestion of clinical dietary treatment for patients with GDM. The experimental data was collected during routine treatment of healthy and GDM pregnant women at Peking University People’s Hospital (see Methods). Two cohorts of 27 patients with GDM and 30 healthy pregnant controls were recruited for this study. All subjects were between 24 and 28 weeks of gestation at the start of the study. The average age of the 30 subjects in the control group was 31.4 years, with an average pre-pregnancy BMI of 21.3 before pregnancy, and an average BMI at sampling of 25. The average age of the 27 patients with GDM was 32.7 years, with an average BMI of 24.1 before pregnancy and 27.04 at sampling. 16S rRNA analysis of stool samples collected twice over a two-week interval represents the microbial communities at the operational taxonomic units (OTUs) level. These sample sets are notated as g(W0) and g(W2) for the GDM subjects and h(W0) and h(W2) for the healthy subjects (Fig. 1a). GDM patients had diet intervention treatment during these two weeks. Blood glucose levels were collected following the OGTT and routine monitoring. In the first sampling, the OGTT was performed on all subjects, and was used to classify the subjects as either healthy or GDM (Fig. 1b). During the second sampling, after two weeks, normal routine blood glucose monitoring was performed on the GDM group alone (Fig. 1b). Of the original cohort, 7 pregnant women completed the second glucose test.

**Fig 1.**
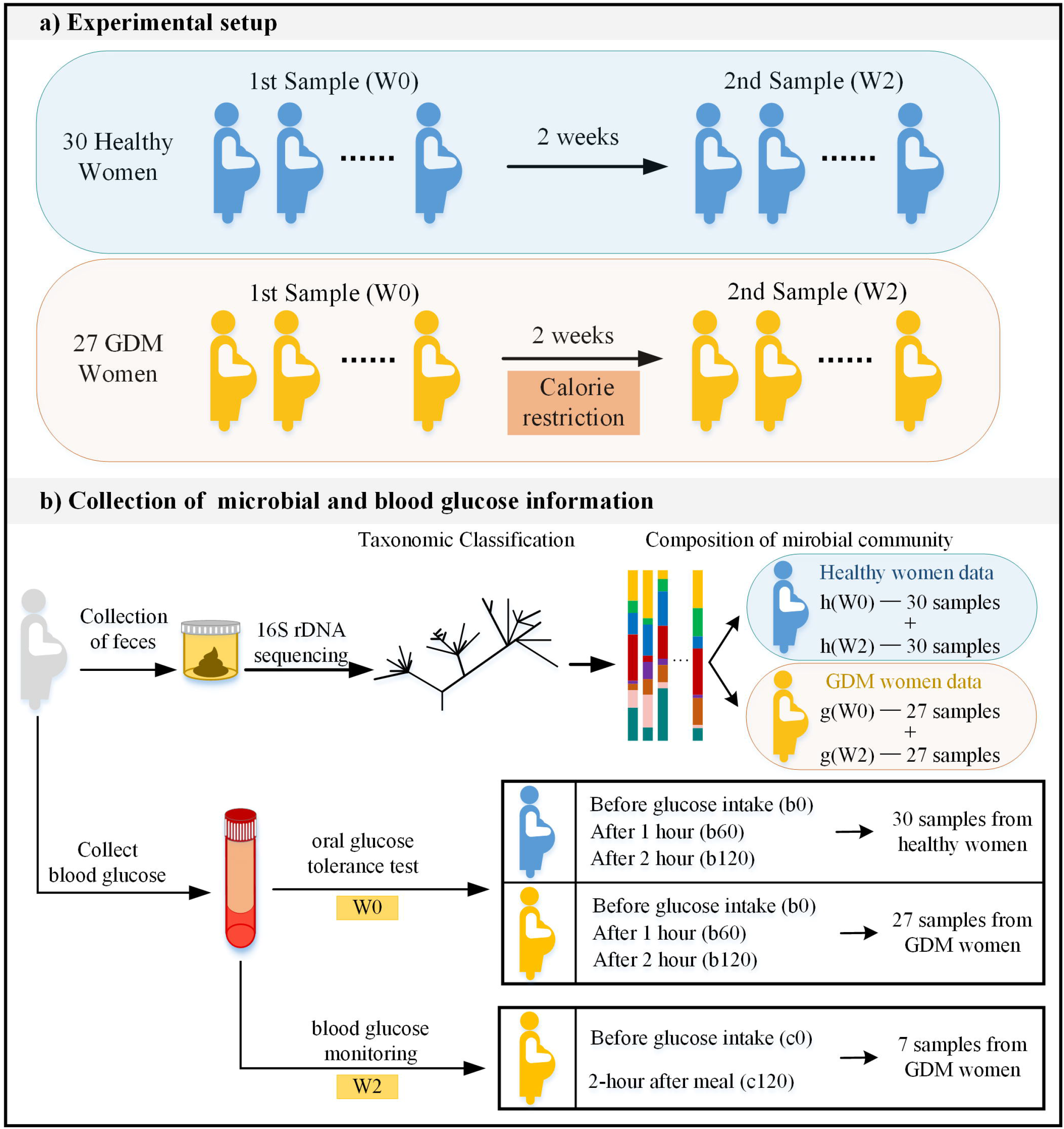
Description of the experimental setup and data collection process. a) Experimental setup - Subjects include 27 GDM patients and 30 healthy pregnant women as the control group. Samples were collected twice for each subject, at interval of two weeks. The GDM patients executed diet intervention for two weeks. b) Collection of microbial and blood glucose information - To collect microbial information, DNA was taken from all subjects through feces, using 16S rDNA sequencing to derive taxonomic classification of microbiome and microbial community. To collect blood glucose information, 75 gram OGTT was performed on all subjects at the first sampling and routine blood glucose monitoring was performed on 7 GDM patients at the second sampling.

OTUs co-expression networks were reconstructed for the four different sample groups. In our study, network analysis includes two types: group analysis and individual analysis (Fig. 2, see Methods section). Group analysis compares the similarity of microbial networks between two groups of subjects (Fig.2a). Individual analysis measures how much the ecological balance of an individual subject is consistent with the ecological balance of the rest of the subjects in the same group. We estimated the individual’s network-impact with a ‘leave-one-out’ procedure (inspired by the method described in [30]). Specifically, we introduced two ways to evaluate the network-impact of an individual sample. First, we compared the network structure reconstructed from samples without the interested sample and with the interested sample (Fig. 2b). Second, we measured the impact of the interested sample with respect to reference samples indirectly (Fig. 2c). The network analyses were also accompanied by traditional microbial community analysis for the purposes of comparison.

**Fig 2.**
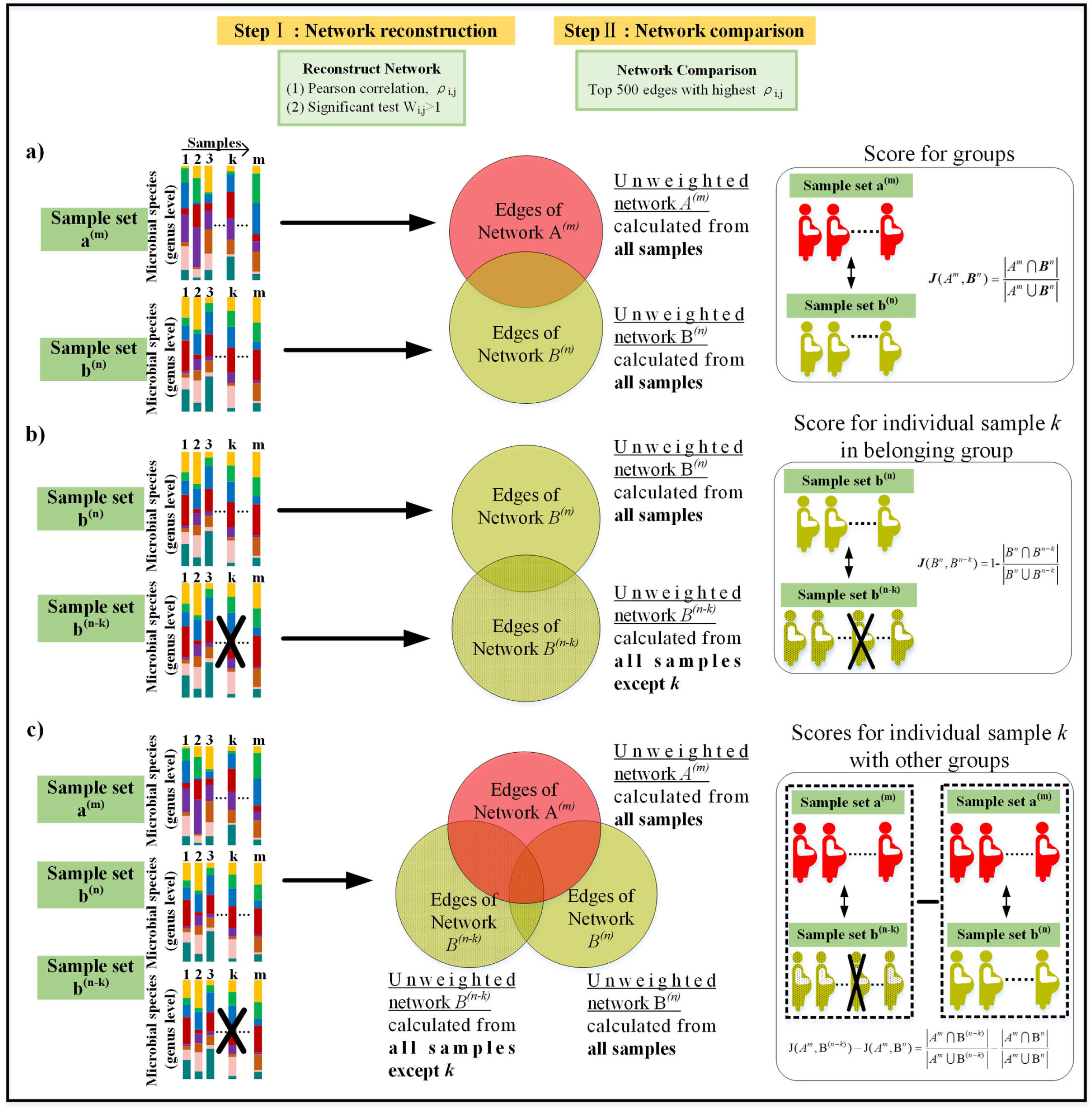
Overview of network reconstruction method and evaluation. For each sample set, a network of pairwise interaction was constructed. Network edges are constructed according to correlation value, and the set of edges were used to evaluate the overlap between networks. a) The method of evaluating the similarity between two different sets. b) The method of evaluating the impact/effect of each individual on the network reconstructed for its group. c) The method of evaluating the impact/effect of each individual on the network reconstructed for the other groups.

### The microbiome composition in patients with GDM and healthy pregnant subjects

After the OTU filtering procedure, 108 OTUs were left in each group (see Methods section). First, we compare the similarity between different groups by community analysis methods. For the beta diversity analysis, the root Jensen–Shannon divergence is used (rJSD) [31] to calculate the dissimilarity of the different sample sets. For the PCoA analysis, we again use the rJSD metric to calculate the distance distribution between the different sets. It is found that the PCoA of the microbial composition of the healthy subjects and subjects with GDM before and after two weeks of diet intervention shows no significant differences among the four groups in the microbiome community structure (Fig. 3a). For each group, we also calculate the beta diversity, measured as the pairwise distances among all samples in the same group (Fig. 3b). The Wilcoxon rank-sum test shows no apparent significant differences between any two groups (P-value>0.05). This implies that it is also difficult to observe the differences in the microbiome community of pregnant women under the OTU scale using the traditional microbiome community analysis method. In addition, we have systematically tested for diet-related changes, i.e., G(W0)/G(W2) comparisons, in all the individual taxa in our data (species taxonomic level). We have found no individual taxa with a significant differential abundance (p-value>0.05 for all taxa, Mann-Whitney U-test with Bonferroni correction for multiple comparisons). Traditional microbiome community analysis methods mainly focus on the differences in microbiome abundance values in individual taxa, but often ignore the interactions between different taxa, which may be capture using network approach.

**Fig 3.**
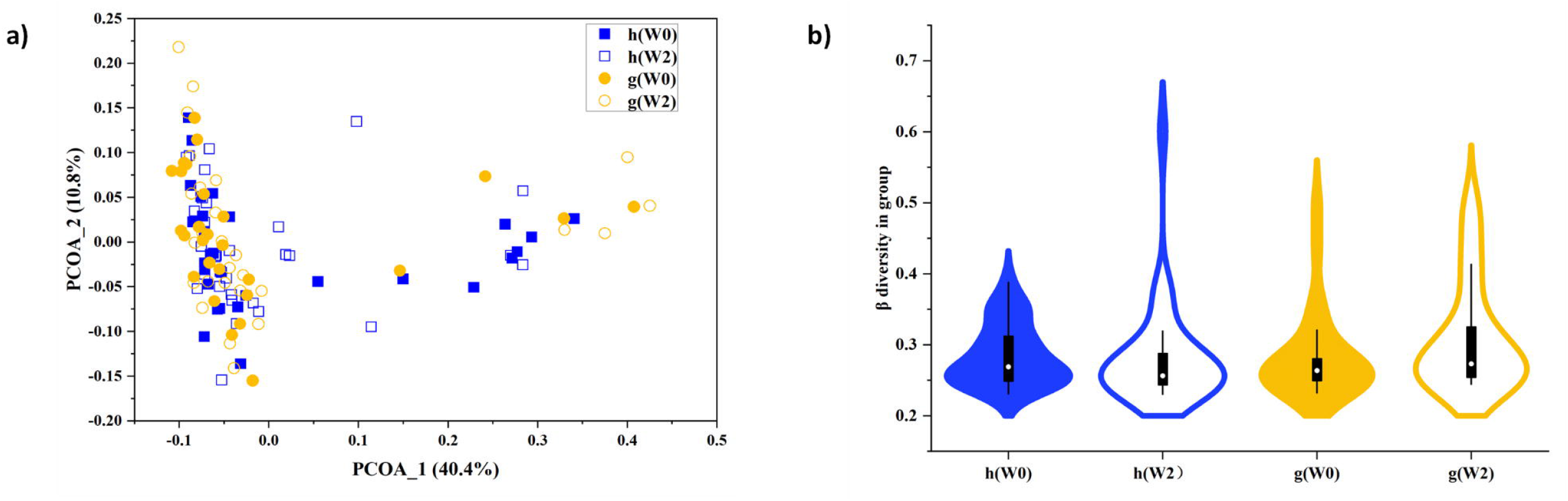
Community analysis of the gut microbiome composition of healthy and GDM patients. a) Principal Coordinates Analysis (PCoA) plot showing four groups of subjects. The horizontal and vertical coordinates are the first two principal components respectively, and the percentages in parentheses are the percentages of variables that can be explained in terms of principal components. b) Violin plot of beta-diversity among subjects within the same group calculated by rJSD distance. The samples show no apparent significant differences between any two groups of them (P-value=0.22, 0.51, 0.38, 0.79, 0.06, 0.15 separately using the Wilcoxon rank-sum test). The number of samples in healthy group (h(W0) and h(W2) is 30 and the number of samples in GDM group (g(W0) and g(W2) is 27).

### The stability of the microbial networks

Next, we use the network analysis methods to analyze the networks’ stability among the different groups. We first calculate the Jaccard similarity between the microbial networks of the healthy group reconstructed from samples collected at W0 and W2 and compare it to the Jaccard similarity calculated between two shuffled networks. This shuffled model represents two independent networks, while preserving the number of links of the original networks (see Methods). The Jaccard similarity of the healthy group is ∼0.2, which is almost 4 times higher compared with the similarity between the shuffled networks (∼0.05) (Fig. 4b). This represents the consistency level of the network after two weeks for the same group, even without any known perturbation (such as diet intervention). Similarly, the Jaccard similarity calculated between GW0 and GW2 (0.185) is also significantly higher compared with the shuffled model, demonstrating its overall level of stability. This level of stability (about 0.19) may reflect the dynamic of the microbiome during pregnancy or the technical inaccuracy of the network reconstruction procedure and represents a baseline for the following analysis. Importantly, the fact that the Jaccard value of the GDM networks before and after the two weeks is lower compared with the healthy networks is inconclusive since it may be associated either to the GDM condition itself or to the diet intervention.

**Fig 4.**
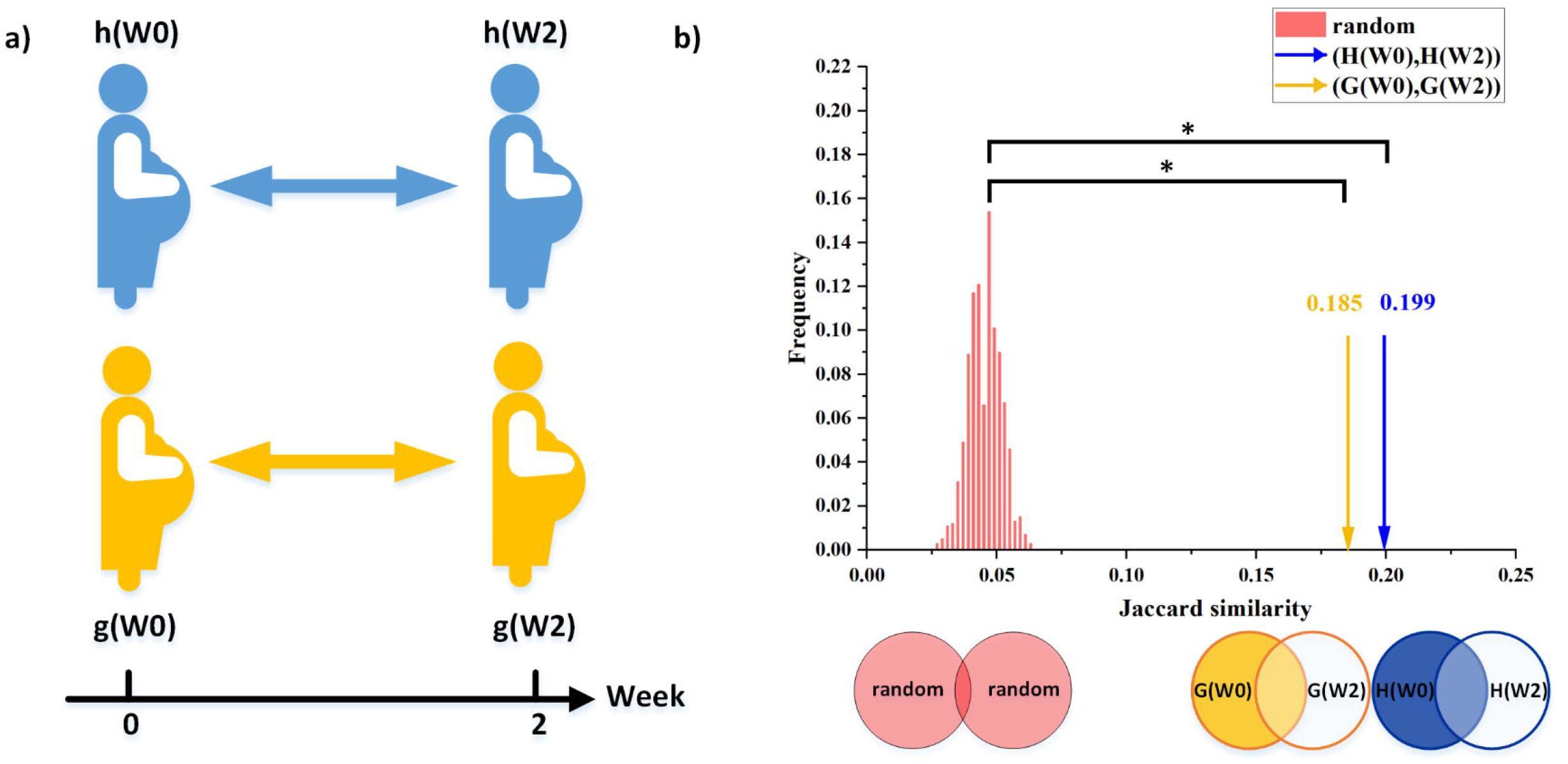
Comparison of the healthy group similarity by network analysis. a) The consistency of the microbiome network of the GDM group and the healthy group after two weeks are evaluated by comparing the similarity of the GDM group and the healthy group to the null model. b) Comparison between the similarity score for the healthy group and GDM group before and after the two week interval and the score of the groups of null models created using a shuffling procedure (see Methods). In the null mode, the edges between the nodes are randomly shuffled, preserving the overall network size. The similarity score for the unshuffled data is marked with yellow arrow and blue arrow, representing Jaccard similarity between H(W0) and H(W2), G(W0) and G(W2), respectively. H(W0), H(W2), represents the network constructed by h(W0) and h(W2), respectively. H(W0), H(W2), represents the network constructed by h(W0) and h(W2), respectively. G(W0), G(W2), represents the network constructed by g(W0) and g(W2), respectively. The significant similarity between the networks calculated for the same subjects after a two weeks interval indicates that they capture a consistent pattern of the inter-species correlations. * indicates *p* < 10^−3^ calculated as the fraction of shuffled realizations with Jaccard value equal or larger than the observed value.

### The effects of diet intervention on GDM patients

We next investigate the effects of diet intervention by analyzing the change in the microbiome community balance level of GDM patients after diet intervention and comparing it to the microbiome of the healthy subjects. Analysis was performed both on the community structure and the co-abundance network level. To reduce biases, we do not directly compare the microbiome community balance of GDM patients and healthy subjects before and after the diet intervention, because the differences in results may be due to diet intervention or disease causes. Instead, in order to make a more effective comparison, we evaluate the change in the GDM microbiome indirectly by measuring its similarity to the healthy microbiome, which serves as a reference group (Fig. 5a). For the network analysis method, network similarity calculation is performed to identify the balanced health of the microbial community in GDM patients during diet intervention. Surprisingly, after two weeks of diet intervention, the similarity between the networks of the GDM patients and the healthy patients is significantly reduced (Fig. 5b P-value<10^-9^ using the Wilcoxon rank-sum test). Besides the network analysis method, we compare the similarity between different groups by β diversity analysis, too. By calculating rJSD distances of microorganism between different groups samples, we found in contrast that there is no significant differences in the distance between the GDM patients and the healthy community (Fig. 5c, P-value>0.02 using Wilcoxon rank-sum test).

**Fig 5.**
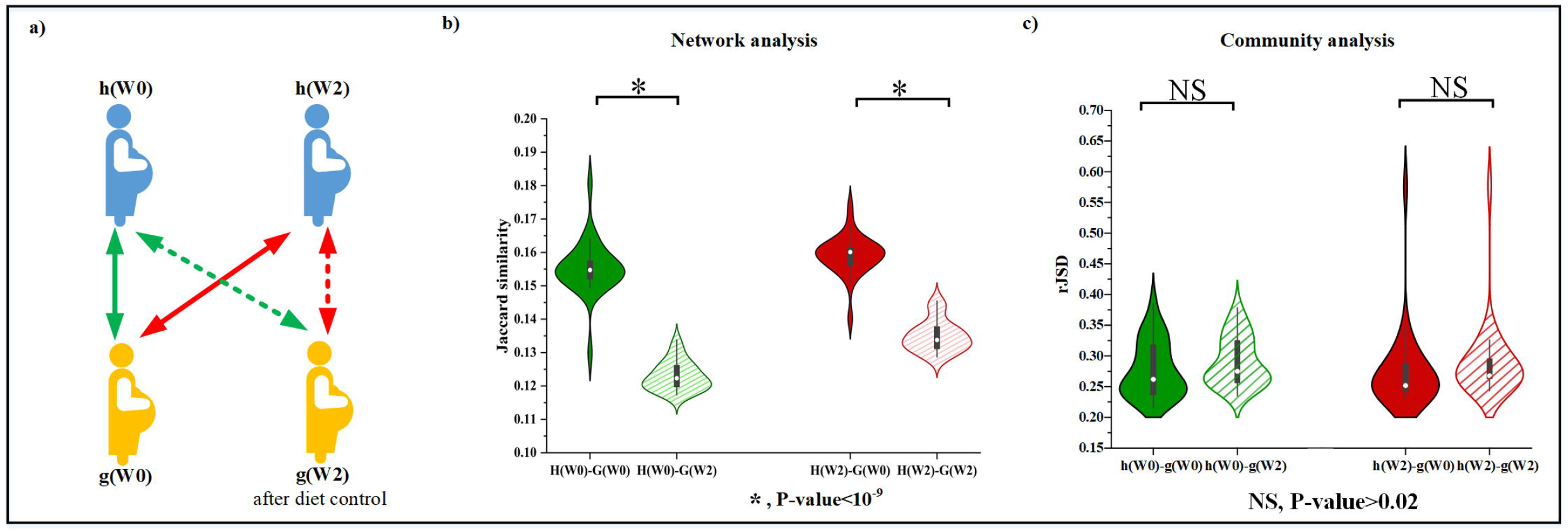
The effects of diet intervention on the gut microbiome as expressed by network and community analysis. a) The effect of the diet intervention on the microbiome of the GDM patients was evaluated indirectly by comparing it to the reference group of the healthy patients. b) Violin plot of the Jaccard similarity between the networks of the GDM group after diet intervention and the healthy group (shadowed areas) was significantly lower than before the diet (filled areas) (P-value=3.72e-10 and 4.37e-10 using Wilcoxon rank-sum test). In addition, the similarity of H(W0)/G(W0) is significantly higher than the similarity of H(W2)/G(W2) (p-value =3.6e-9 using Wilcoxon rank-sum test). c) Violin plot of the dissimilarity between different groups by community analysis method. Each value represents the average distance (rJSD) calculated between each of the GDM samples and the samples of the reference group (healthy). The samples show only minor variability (P-value=0.1323 and 0.0214 using Wilcoxon rank-sum test).

These results demonstrate that the microbial communities are altered during the two-week diet intervention period. This change is not captured by traditional beta-diversity analysis or by distance measures but is instead reflected in the ecological networks. Moreover, the direction of the change observed by our ‘indirect comparison’ was counterintuitive. While diet intervention is clinically beneficial to the GDM patients, the underlying ecology of the patients’ microbiome was not ‘healthier’, i.e., it was less similar to the healthy group. In the future we hope to discern whether these changes in the microbial co-abundance correlation have direct casual relations to the health benefits of diet intervention in the GDM patients.

### Associations between ecology of microbial network and abnormal glucose patterns

Finally, we study the relationship between the microbiome of individual GDM patients and their blood glucose measures. By analyzing individual microbiomes balance in GDM group, we hope to find out the specificity of individuals in GDM patients and analyze whether this specificity is related to changes in blood glucose, so as to provide support for personalized medicine. We apply microbial network analysis method to evaluate the differences in microbiome balance between each GDM individual and others in GDM patients by calculating the network similarity. While the microbial networks represent the group-average relationships between the microbes, each subject has a unique individual signature of microbial co-abundance relation, and its specific networks can reliably describe individual specific disease states [30]. Using the "individualized network analysis" methodology (see Methods), we analyze the microbial samples of individual subjects based on their microbial network and compare it to the patterns of blood glucose levels from the OGTT and routine blood glucose monitoring. To analyze the pattern of microbial co-abundance community balance in individual subjects, we first perform a ‘leave-one-out’ procedure which compares between the networks calculated without each patient and the ones calculated with it (Fig. 6a, see Methods). We directly measure the changes in the network structure reconstructed from a cohort of samples after removing the individual sample *k* of interest before diet intervention. Figure 6a shows that the Jaccard distance of patients number g2, g13 and g23 are significantly higher compared with the other patients using the Wilcoxon Signed-Rank Test (P-value= 1.18E-05(P-value= 2.35E-05, P-value= 8.30E-06, respectively), suggesting that the community balance of these individuals differ substantially from the others in the group.

**Fig 6.**
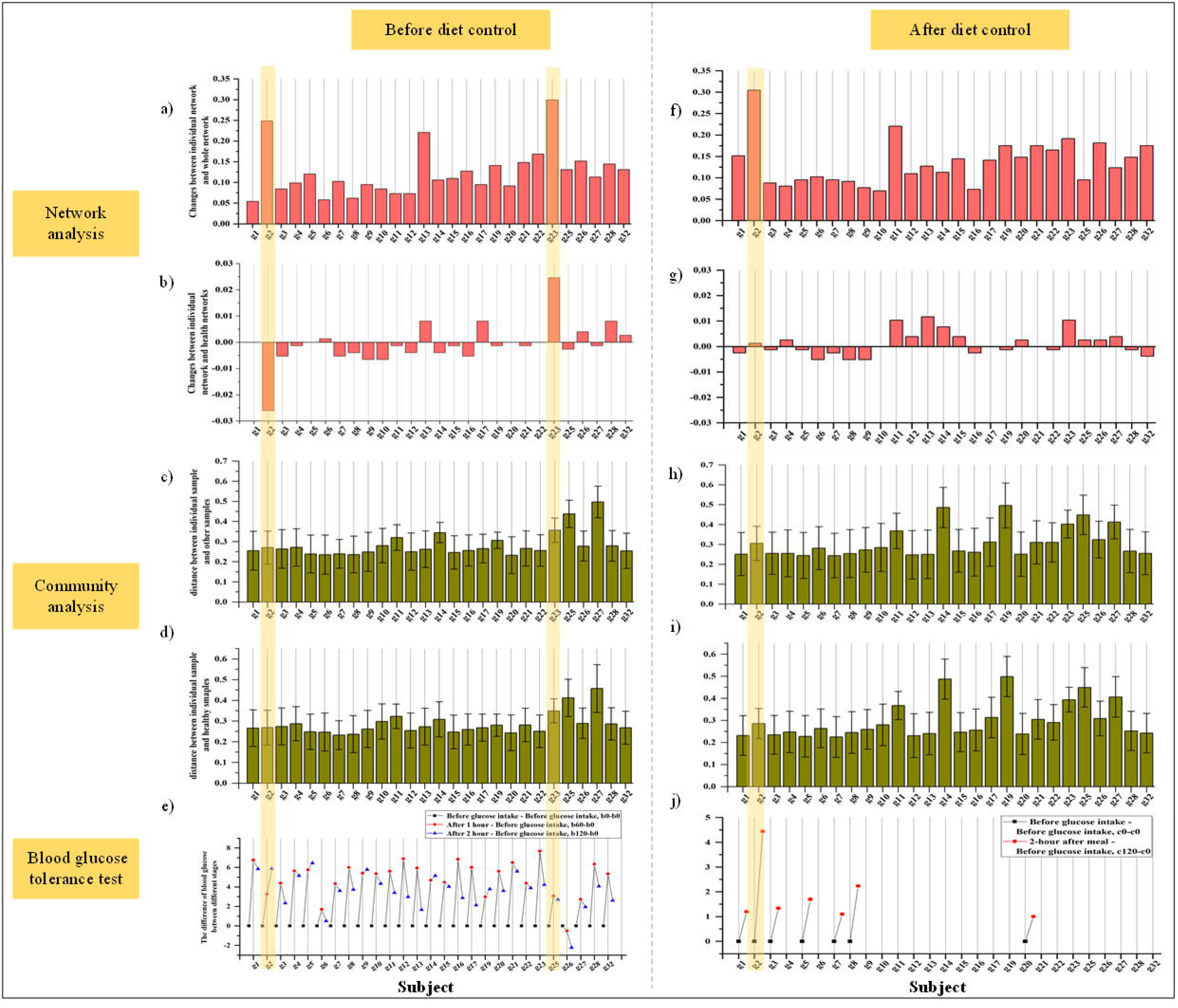
Analysis of the relationship between microbial system and blood glucose levels. a) Evaluation of the changes between individual network of each GDM patient and the whole network of all GDM patients by Jaccard dissimilarity score before diet intervention. b) Evaluation of the impact of sample *k* of GDM patients on the network of all GDM patients by measuring its change with respect to the healthy women by Jaccard distance score before diet intervention. c) Evaluation of the dissimilarity between each individual GDM patient to other GDM patients using the root Jensen– Shannon divergence (rJSD) before diet intervention. d) Evaluation of the dissimilarity between individual GDM patient to healthy women using the rJSD before diet intervention. e) OGTT blood glucose information collected by all GDM patients before diet intervention. f) Evaluation of the changes between individual network of each GDM patient and the whole network of all GDM patients by Jaccard dissimilarity score after diet intervention. g) Evaluation of the impact of sample k of GDM patients on the network of all GDM patients by measuring its change with respect to the healthy women by Jaccard distance score after diet intervention. h) Evaluation of the dissimilarity between each individual GDM patient to other GDM patients using the root Jensen–Shannon divergence (rJSD) after diet intervention. i) Evaluation of the dissimilarity between individual GDM patients to healthy women using the rJSD after diet intervention. j) Routine blood glucose monitoring information collected by 7 GDM patients after diet intervention. The shadow box corresponds to the subject with abnormal blood glucose regulation whose impact is significantly higher than others according to the microbiome network.

Based on network analysis method, we evaluate the differences in microbiome community balance between the GDM individual and the healthy group. Specifically, when one individual from the GDM group is dropped out, we calculate what extent the similarity level between the networks of the GDM and the healthy groups change (Fig. 6b). The equation in Fig. 2c is used to analyze the network impact of each GDM patients before diet intervention. We choose the healthy group before diet intervention as the reference cohort and evaluate the impact of sample k in GDM patients on the network of its cohort by measuring its change compared to the healthy group before diet intervention. Fig. 6b shows that patients g2 and g23 also exhibit a clear individualized impact using the Wilcoxon Signed-Rank Test (P-value= 3.09E-07, P-value= 8.29E-06, respectively).

The differences in microbiome community balance of g2 and g23 can correspond to abnormalities in the blood glucose levels before diet intervention. The blood glucose data measured by OGTT has a repeated pattern across the subjects (Fig. 6e). Fasting blood glucose (b0) is usually the lowest of all other measurements. The blood glucose level after drinking glucose solution for 1h (b60) is the highest, and after 2h (b120), it decreases due to human body regulation, but is still higher than the fasting blood sugar level. Even though all GDM patients have blood sugar values that are higher than the healthy population, some of which seem to stand out within the group, indicating uniquely abnormal blood glucose regulation. For example, for patient number g2, the blood glucose level of patient number g2 gradually increases over time, and the 1-hour blood glucose level increase of patient g23 is exceptionally higher compared with the other patients.

Correspondingly, we evaluate the differences in microbiome community balance between the GDM individual and others of patients/healthy after diet intervention (Fig. 6f and 6g). Specifically, patient g2 stood out in the analysis. When we evaluate changes in the similarity level between the networks of the GDM and the healthy group while dropping out g2 from the GDM, we found that the change between microbial networks of the g2 and healthy groups networks after diet intervention was not as large as they were before diet intervention (Fig. 6b). Although the similarity between the microbial network of g2 and that of the healthy group is the same as the similarity between the complete group network and the healthy group at this time-point, the individual microbial network of patient g2 has a very low similarity to the complete group network.

After diet intervention, the relationship between microbiome community balance and abnormal blood glucose was still seen. Blood glucose data was measured using routine blood glucose monitoring. In total, 7 patients reported self-monitoring of blood glucose levels (Fig. 6j). We find that the 2-hour postprandial of g2 is higher than other patients. The abnormalities of g2’s blood glucose correlates with the observed phenomena in the microbiome network. This implies that abnormal blood glucose regulation in GDM patients is related to the interactions/ecology of the microbial network. Simply put, the individual network method analysis suggests that the blood glucose regulation level of GDM patients is partially related to its microbial composition.

We find rare similar evidence from traditional microbial community analysis method. For each GDM subject, the β diversity analysis is performed by calculating rJSD distances to compare the dissimilarity between this GDM subject and the other GDM subjects (Fig. 6c and h). Additionally, we perform the β diversity analysis by calculating rJSD distances to compare the dissimilarity between this GDM subject and the all healthy subjects (Fig. 6d and i). We found that when applying the microbial community analysis method, there is no apparent significant relationship between the blood glucose regulation level of GDM patients and its microbial. By calculating the differences between the microbial community structure of a specific individual and other individuals’ microbial communities’ structure both before and after diet intervention, we find that subjects with large differences do not correlate with those with abnormal blood glucose. For example, we find patient g2 is abnormal in blood glucose level. But the microbial community structure of g2 was not significantly different before diet control from that of other individuals using the Wilcoxon Signed-Rank Test (P-value= 0.949). Similarly, when comparing the microbial community structure of a specific individual with that of the healthy group, a similar conclusion is found: no relationship between blood glucose levels and microbial composition is observed using traditional microbial community analysis methods. The microbial community structure of g2 after diet control was not significantly different from that of other individuals using the Wilcoxon Signed-Rank Test (P-value= 0.665).

However, the anomaly of the g2 patient does not represent a typical microbiome pattern in the analyzed patients. The microbiomes and microbial networks of patients g5, g9, g14 and g19, which exhibit a similar, but less pronounced, blood glucose anomaly, are not more similar to g2 than the other patients. Further research on larger cohorts is required to test whether there is a common mechanism that links blood glucose and the microbiome.

## Discussion

Personalized medicine require more precise identification of each individual. In this work, we characterize the microbiome from its network interaction in the individualized level. We analyze the microbiome of patients with GDM and healthy subjects through the lens of network analysis. For the implementation of personalized health management of GDM patients, it is very important to explore individual differences from the perspectives of physiological indicators and living habits. In individual network analysis we found that abnormal glucose regulation is associated with large network deviations, which may lead to the development of individualized microbiome-based therapies in the future. Previous work that analyzed the composition of intestinal bacterial flora at two time-points of subjects under traditional microbiome analysis method concluded that overall bacteria gathered in response to diabetes status, rather than diet intervention. Short-term diet management plays a role in the process of GDM by affecting specific taxa. Short-term dietary management is not an alternative pattern for gut microbial [32]. Here, in contrast, network analysis enabled us to find changes in the dynamic interactions among microorganisms in the community balance of the microbiome that are undetected with traditional approaches.

Our goal is to study the *network similarity* between groups, a concept which is fundamentally different from the standard *community similarity*. From the perspective of community similarity, we see no significant difference between the microbiomes of the healthy and the GDM groups, both before and after diet intervention. However, from the perspective of the microbial networks, the diet intervention has a clear effect. Surprisingly, after the diet the microbial networks of the GDM group become less similar to the healthy compared with their state before the diet.

We conclude that diet intervention is a treatment that could help GDM to balance their blood glucose to control the disease but does not necessarily benefit the microbial ecological balance. In fact, some treatments do break the balance of the microbial community in order to treat patients. For example, the use of antibiotics, which can speed up treatment, should be avoided to prevent affecting local microbiota, as it may contribute to obesity and type 1 diabetes [32–38].

Besides, our research emphasis is to analyze the individual patient. A safer and more effective treatment can be achieved by personalizing the general recommendations [39]. Our study find that abnormal microbiome balance is associated with abnormal glucose regulation. Our method can analyze individual patients through individual network analysis to evaluate the degree of abnormal glucose regulation, which reflects GDM patients’ ability to regulate blood sugar after sugar intake. Therefore, according to the different situation of each patient, we could potentially implement more effective and reasonable diet intervention strategy or other treatment that not only rely on the patient’s body indicators such as height and weight, but also consider the patient’s individual blood glucose regulation level. Based on this study, we can further and better carry out individualized precision medicine for GDM. For example, with a clearer description of the expected effects of diet intervention on GDM patients, we might be able to monitor new patients by comparing their microbiome to representative cohort and checking whether their microbiome evolution trajectory follows the norm.

## Materials and Methods

### Subjects and sampling description

Samples were gathered at Peking University People’s Hospital during 2017 from 27 patients with GDM and 30 healthy pregnant subjects (control group), who were selected according to their matched age and gestation period. Make sure all subjects are with no antibiotic selection and with no concurrent 83 diseases during the 3 months before sample collection. For each subject, microbial and blood glucose samples were collected twice, in two-week intervals. For patients with GDM, calorie restriction was implemented through daily diet intervention during these two weeks, as described below. For the control group, no calorie control was implemented.

Fasting 75 g OGTT is chosen to diagnose the pregnant subjects between 24 and 28 weeks gestation, which is the primary diagnostic method of GDM. The test involved drinking a solution containing 75g glucose, and drawing blood to check glucose levels at 0h and after 1h and 2h. GDM is diagnosed if one or more level(s) elevated. The thresholds for OGTT are 5.1 mmol/L at 0 hour, 10.0 mmol/L at 1 hour and 8.5 mmol/L at 2 hours during OGTT, respectively. This thresholds is suggested by the International Association of the Diabetes and Pregnancy Study Groups in 2011.

### Diet intervention strategy

The macronutrients (protein, fat and carbohydrate) and caloric consumption of GDM patients were estimated during the two weeks diet intervention in consultation with a nutritionist. Participants were deemed to have complied with the given dietary recommendations when all of the criteria below were met: 35–45% in total energy is carbohydrates, low glycemic index carbohydrates and 20% in total energy is simple carbohydrates. 18–20% in total energy is proteins and 35% in total energy is fats. at least 20–25 g/day for fiber intake, and make sure no alcohol consumption. The recommended daily calories are divided into smaller, multiple meals to protect patients from ketonuria and acidosis because it often occurs due to prolonged fasting. Besides, The nutritionist was contacting with subjects with GDM continuously, through telephone contact every week, to keep them updated on their nutritional status as the study progressed. Besides, the nutritionist instructed patients to monitor blood glucose by themselves at least 4 times a day by finger puncture capillary blood glucose test. To avoid the the gut microbiota composition to be effected by prebiotics/probiotics use, general recommendations were as implemented for the healthy pregnant subjects, making sure no spicy foods and no yogurt intake.

### Microbial data extraction method

#### DNA Extraction & OTU analysis

Stool samples were frozen as soon as possible after being collected and stored at −80 °C until DNA extraction was performed as described in [40]. Base on the manufacturer’s instructions, 200 mg was extracted from each feces sample for DNA extraction by the QIAamp DNA stool Mini kit (Qiagen, Germany). 515F (5’-GTGCCAGCMGCCGCGGTAA -3’) and 806R (5’-GGACTACHVGGGTWTCTAAT -3’) are used to amplify the V4 region of the 16S rRNA. Each appropriate sized PCR product was purified and then use the HiSeq 2500 genome analyzer (Illumina HiSeq 2500) to perform the 250-bp nucleotide paired-end sequencing. High-quality trimmed reads were aggregated into OTUs by MOTHUR [41], and the recognition rate was 97%. To make sure the phylogeny of the OTUs, using the Greengenes database to BLAST search the longest sequence from each OTU [42] to obtain full-length 16S rRNA gene sequences with well-annotated full-length.

#### Data pre-processing

Our initial dataset contained 57 subjects with 813 unique OTUs identified. In order to avoid artifactual/spurious associations between non-correlated and low-abundant microbial members in a community, OTUs that were found in less than 10 instances or were found in less than 10% of all subjects were filtered out. The remaining OTUs were used to reconstruct the co-abundance networks. Considering that there will be a large variability in the microbial abundance values, in order to make the calculation results more reliable, the microbial abundance data of each subject is normalized to make the sum of the microbial abundances of each subject equal to 1. Then, the samples was divided into four parts for analysis according to subject type and sampling time, including first sampling data of 30 healthy pregnant women, second sampling data of 30 healthy pregnant women, first sampling data of 27 GDM patients and second sampling data of 27 GDM patients.

### Network analysis method

#### Network reconstruction principle

For the four sample groups of different states (h(W0), h(W2), g(W0), g(W2)), we reconstructed the OTUs co-expression binary networks. Each node in the networks represented a single OTU. The edges of the network corresponded to significant correlations between pairs of OTUs. The following processes were applied to reconstruct the networks: (1) For each group, the Pearson correlation for all pairs of OTUs was calculated; (2) non-significant correlations were filtered out using a Z-score test. For each pair of OTU sequences, the samples were randomly shuffled 1000 times and the Pearson correlation coefficient calculated. Then, the Z-score, *W,* was calculated according to the following formula:

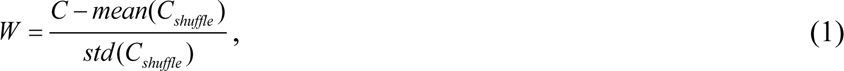

where *C* is the Pearson coefficient of the non-shuffled data, *mean(C_shuffle_)* is the average value of Pearson coefficient of the shuffled data and *std(C_shuffle_)* is the standard deviation value of the Pearson coefficient of the shuffled data. Larger W value means that the correlation is more significant. A value of W < 1 was considered a non-significant correlation and filtered out; (3) For each network, a fixed number of 500 edges were defined as the OTU pairs with the highest Pearson correlation values. The reasons and necessity of fixing the size of network are elaborated in the supplementary information (S1 Fig, S2 Fig and S3 Fig). This step was necessary for eliminating the possible bias of the number of edges when comparing the structural similarity between different networks.

#### Group network analysis

To compare between two groups, networks were reconstructed using all the samples of each group. The similarity between the networks was defined as the overlap between the set of edges, according to the Jaccard index:

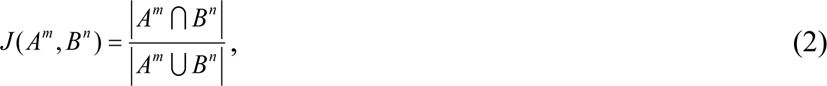

where *A* and *B* are two different sample sets, *m* and *n* are the number of subjects in each sample set. *A^m^* and *B^n^* represent the set of edges of the two networks, respectively.

#### Individualized network-impact

Inspired by the LIONESS method for inference of single-cell gene regulatory networks [43], our network reconstruction method analyzed the *network-impact* of individual GDM patient samples. However, unlike the LIONESS method, our method did not aim to infer the network entirely, but to simply evaluate the impact of a single sample in general. In order to measure how much the ecological balance of individual subject *k* is consistent with the ecological balance of the rest of the subjects in the same group, its *network-impact* was estimated with a ‘leave-one-out’ procedure (inspired by the method described in [44]). Specifically, two ways to evaluate the network-impact of an individual sample, *k,* were introduced.

The first way directly measured the change in the network structure reconstructed from a cohort of samples after removing the individual sample of interest. The Jaccard dissimilarity score was calculated,

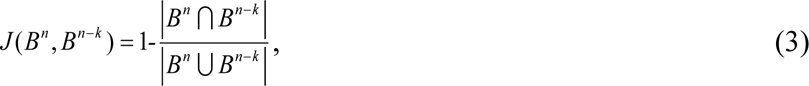

where *B^n^* represents the network that was reconstructed with all samples *and B^n-k^* represents the network that was reconstructed without sample *k.* Low dissimilarity indicated that the balance between species abundance of sample *k* tend to follow the same correlation pattern of the entire group, while high dissimilarity suggested that sample *k* follows a unique correlation pattern.

According to Eq. (3), the larger the Jaccard distance, the lower the similarity between the network without the sample *k* and the network with the sample *k*. This suggests that this sample *k* made a significant difference in all samples. When the Jaccard distance is 1, it means that the network without the sample *k* is completely different from the network with the sample *k*. Alternatively, when the Jaccard distance is 0, it means that the network without the sample *k* is exactly the same as the network with the sample *k*.

The second way is an indirect evaluation of the impact of sample k on the network of its cohort by measuring its change with respect to a reference cohort. The change in the Jaccard distance score was calculated

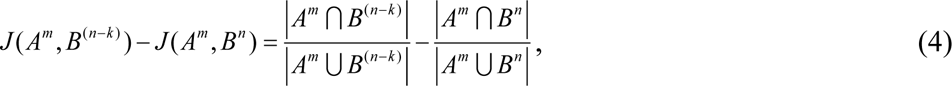

where *A^m^* represents a network that was reconstructed from the reference cohort. A small (large) change in the distance between networks A and B after removing sample *k* indicates that *k*’s abundance profile follows a similar (different) correlation pattern to its cohort.

Here, it can be seen from Eq. (4) that when the Jaccard distance value is positive (negative), it indicates that network B after removing sample k is more similar (different) to the reference cohort A, and the sample k is the person who is more different (similar) with the reference cohort than the others.

#### Data and network shuffling processes

*Data shuffling*: In the network reconstruction process, the significance of each edge was estimated by comparing its associated Pearson correlation value to a set of values calculated for shuffled abundance profiles. Each shuffled abundance profile was reconstructed using a Monte Carlo procedure, by randomly assigning a value for each OTU from the empirical abundance distribution of the same OTU, independently. The shuffled profiles preserve the original relative frequencies of the OTUs while removing any correlations among them. *Network shuffling*: The distance values between networks were compared to distances calculated between shuffled networks, reconstructed with the same number of nodes but with random reassignment of the 500 edges.

### Community analysis method

In addition to network analysis methods, the microbiome community composition in different groups were compared and analyzed by calculating the β diversity according OTU table. We calculate the dissimilarity of different sample sets by using the root Jensen–Shannon divergence (rJSD) measure [31] to compare the difference between different groups. The root Jensen-Sahnnon divergence (rJSD) is defined as

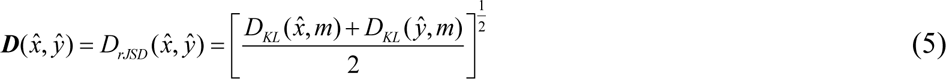

where *x̂* and *ŷ* are renormalized the relative abundances of only the shared species (set S). 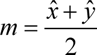 and 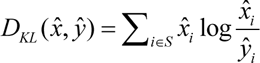 is the Kullback–Leibler divergence between *x̂* and *ŷ*.

## Data availability

The original datasets for this study can be found in the Genome Sequence Archive (https://ngdc.cncb.ac.cn/gsa/), the accession code is: CRA004782. The studied OTU table could be found on GitHub at (https://github.com/YimengLiu9425/code/tree/master).

## Code availability

The Python code used in this study can be found on GitHub (https://github.com/YimengLiu9425/code/tree/master).

## Acknowledgements

This work was supported by the Peking University People’s Hospital Scientific Research Development Funds(grant no.RDY2019-29), the National Natural Science Foundation of China (grant 32070116) and Maternal and Infant Nutrition & Care Research Fund of the Institute of Nutrition and Nursing of Biostime (grant no. 2015-Z-20). A.B. thanks the German-Israeli Foundation for Scientific Research and Development (grant No. I-1523-500.15/2021), the Israel Science Foundation (grant No. 1258/21), and the Azrieli Foundation for supporting this research.

## Authors contribution

N.W. performed the data collection. X.Z. and N.W. performed sequencing management. Y.L. and A.B. developed the methodology and analyzed the data. Y.L., G.A., D.L. and A.B. wrote the manuscript. All authors discussed the results and reviewed the manuscript.

## Ethics declarations

The authors declare no competing interests.

This study was approved by the Conjoint Health Research Ethics Board of Peking University 74 People’s Hospital, and informed consent forms were signed by all of the subjects in this study. All experiments were performed in accordance with the approved 76 guidelines and regulations.

## Inclusion and diversity statement

We support inclusive, diverse, and equitable conduct of research

## Supporting information

**S1 Fig. The size of the microbial networks of the healthy group and GDM patients before and after diet interventions under different W threshold.** The corresponding network size of different groups is different though the threshold is fixed.

**S2 Fig. Jaccard similarity between the GDM group and healthy group with unfixed network size for different threshold values, W.** The green curve shows the GDM group compared with the healthy group two weeks earlier, and the red curve shows the GDM group compared with the healthy group two weeks later. The solid dots indicate the comparison between the GDM group and the healthy group before the dietary intervention, and the hollow dots indicate the comparison between the GDM group and the healthy group after the dietary intervention. The dark curve is the real data result, and the light curve is the shuffled network result.

**S3 Fig. Violin plot of Jaccard similarity between the GDM group and healthy group with fixed network size under different fix number.** When different number of links are fixed, the pattern is still stable in most cases.

