## Supplementary text and figures for "Individualized network analysis reveals link between the gut microbiome, diet intervention and Gestational Diabetes Mellitus"

### Supplementary Notes

***Network analysis details***

In the network analysis, we choose the threshold *W* as 1. When W=1, the network size before and after two weeks in the healthy group and before and after two weeks of diet intervention in the GDM group were 1582,1622,1486 and 1645, respectively. The size of the microbial networks of the healthy group and GDM patients before and after diet interventions were different under different W threshold, as shown in Extended Data Figure 1. Although the threshold is fixed, the corresponding network size of different groups is also different. Therefore, it is necessary to consider whether the size of the network will affect the calculation of network similarity.

In Extended Data Figure 2, the Jaccard similarity of the network is calculated with the size of network unfixed under different threshold W. The dark red and dark green curves show that in most cases, the similarity between the GDM microbiome and the healthy microbiome decreases after diet control. In particular, the light green and light red curves are the Jaccard similarity of shuffled network when the size of network is unfixed under different thresholds W. It is found that when W increases, the network size becomes smaller, and the Jaccard similarity of shuffled network also becomes smaller. If the result of the shuffled network changes with the size of network, it means that the network size will affect the calculation result. Therefore, in order to compare the result of Jaccard similarity between networks more fairly, we choose to fix the size of network.

In this paper, the top 500 most correlated links in the fixed network are selected.

We also show the results for the top 100/300/500/700/900/1100 most correlated links in the network separately, in Extended Data Figure 3. When the number of fixed links is moderate or small, the result are relatively stable. When the number of fixed links was large (e.g., 1100 links), the results were biased when compared to the healthy group microbiome after the dietary intervention. The reason may be that the fixed links are doped with some unstable links because the number of fixed edges is large.

### Supplementary Figures

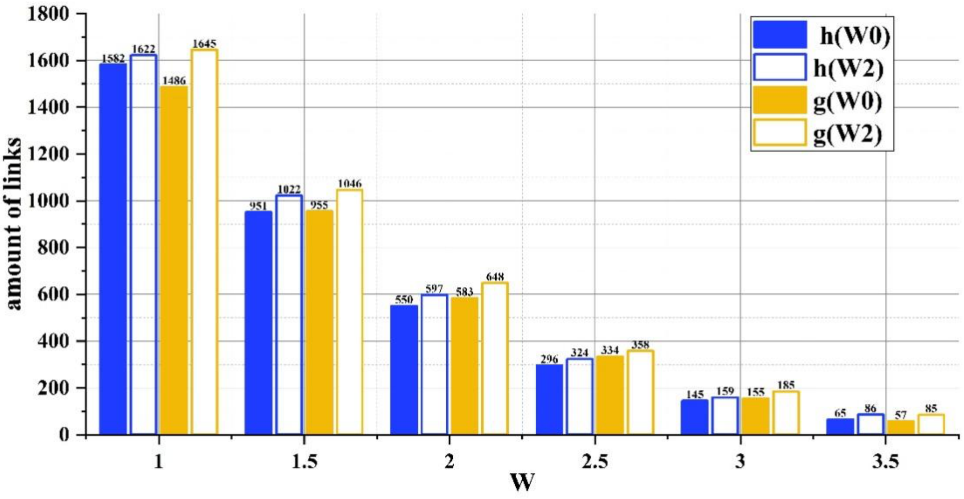


**S1 Figure. The size of the microbial networks of the healthy group and GDM patients before and after diet interventions under different W threshold.** The corresponding network size of different groups is different though the threshold is fixed.

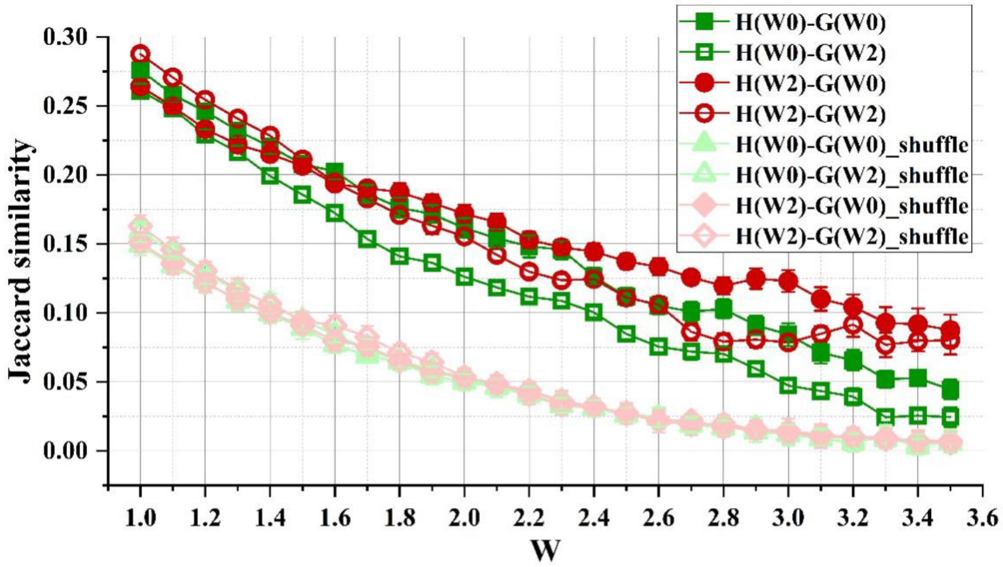


**S2 Figure. Jaccard similarity between the GDM group and healthy group with unfixed network size for different threshold values, W.** The green curve shows the GDM group compared with the healthy group two weeks earlier, and the red curve shows the GDM group compared with the healthy group two weeks later. The solid dots indicate the comparison between the GDM group and the healthy group before the dietary intervention, and the hollow dots indicate the comparison between the GDM group and the healthy group after the dietary intervention. The dark curve is the real data result, and the light curve is the shuffled network result.

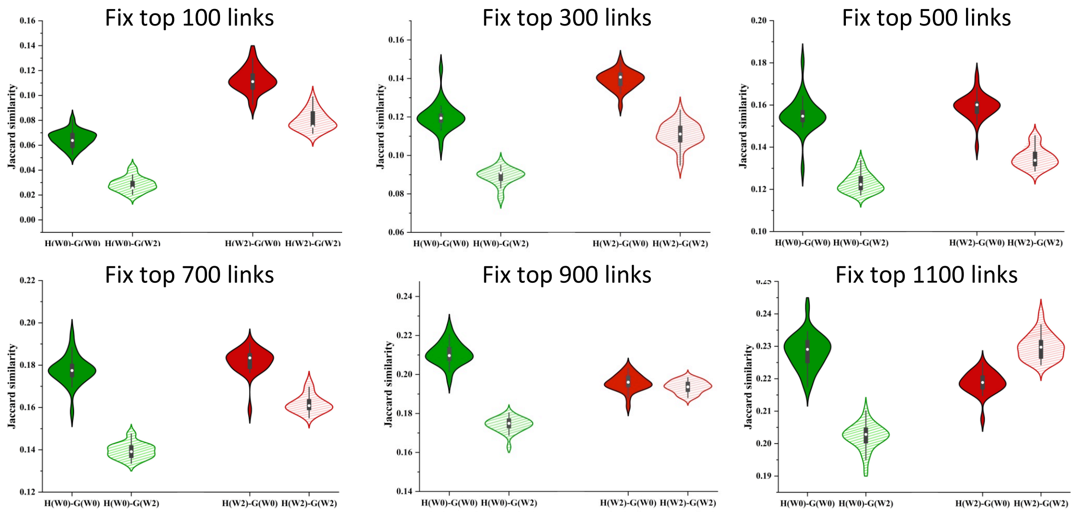


**S3 Figure. Violin plot of Jaccard similarity between the GDM group and healthy group with fixed network size under different fix number.** When different number of links are fixed, the pattern is still stable in most cases.
